# Protective effects of heat shock protein 70 induction against global warming by using oriental bezoar and ginseng

**DOI:** 10.64898/2026.09.08.750282

**Authors:** Eiji Inoue, Tomoe Tsubonoya, Yasuharu Shimizu, Haruhisa Kawasaki, Norio Ishida

## Abstract

The interest in compounds that protect against heat stress-induced damage has been heightened due to world global warming. We found the protective effects of a Japanese natural drug named BG, containing oriental bezoar and ginseng, against heat stress in *Drosophila*. BG suppressed the heat-induced shortened lifespan and reduced fertility in *Drosophila*. Interestingly, the protective effects of BG against heat stress were abolished in heat shock protein 70 (HSP70) mutant flies. To see the protective effects in humans, we applied BG on the cytotoxicity in heat-stressed human hepatic cell line, HepG2. BG suppressed heat stress-induced cytotoxicity at 43°C, and increased HSP70 and heat shock factor 1 (HSF1) mRNA expression in HepG2 cells. These findings indicate that BG protects against heat stress-induced damage via the HSF1/HSP70 pathway and has potential as a therapeutic agent for heat stress-induced disorders, including heatstroke even in human.

## Main Text

Heat stress is a critical environmental factor that disrupts cellular homeostasis and impairs physiological function. Excessive heat exposure induces oxidative stress and protein denaturation, leading to cellular damage, tissue dysfunction, and impaired physiological function (*1*). Heatstroke is the most severe form of heat-related disorder and causes multiple organ dysfunction and death (*2, 3*). The recent global warming trend has heightened the problem influenced by heat stress, there have been many research studies to find novel compounds that protect against heat stress-induced damage (*2–5*).

Among the cellular defense systems against heat stress, heat shock proteins (HSPs) play a central role. Heat shock protein 70 (HSP70) is one of the most important molecular chaperones involved in maintaining protein homeostasis by preventing protein aggregation and promoting refolding of damaged proteins (*6, 7*). In addition to its chaperone activity, HSP70 suppresses apoptosis, stabilizes mitochondrial function, and attenuates oxidative stress-induced cellular damage (*8–10*). Expression of HSP70 is primarily regulated by heat shock factor 1 (HSF1), a transcription factor that is activated upon heat stress by dissociating the binding to HSP90 or HSP70, creating the HSF1/HSP70 pathway (*11*). Activation of the HSF1/HSP70 pathway enhances cellular tolerance to heat stress and protects cells and organisms (*6, 9, 11*). Therefore, compounds that induce HSP70 are considered useful for protection against heat stress-induced disorders, including heatstroke.

BG is a Japanese natural drug containing oriental bezoar and ginseng. Oriental bezoar is a gallstone formed in the gall sac of *Bos taurus* Linné var. *domesticus* Gmelin (*Bovidae*), and ginseng is the root of *Panax ginseng* C.A. Meyer. Oriental bezoar has been described in the Chinese classical medical text Beiji qianjin yaofang as an agent that “eliminates heat” and has traditionally been applied for the treatment of heatstroke (*12*); however, sufficient molecular evidence supporting its efficacy has not yet been established. Ginseng has been widely reported to possess antioxidative, anti-inflammatory, anti-fatigue, and cytoprotective properties (*13*). Previous studies have demonstrated that ginseng ameliorated heat stress-induced reproductive dysfunction in Japanese quails, accompanied by modulation of HSP70 (*14*). Interestingly, the classical text Leigong Yaodui describes that “ginseng serves as an assistant to oriental bezoar” (*15*), implying that ginseng and oriental bezoar may act additively or synergistically. In this study, we investigated the protective effects of BG against heat stress in *Drosophila* and the human hepatic cell line (HepG2), focusing on the involvement of the HSF1/HSP70 pathway.

## Results

### BG suppressed shortening of survival and reduction of fertility in heat-stressed Drosophila

To evaluate the protective effects of BG against heat stress, survival and fertility were examined in heat-stressed flies. Heat stress (30 °C) caused shortened survival (Fig. 1A) and reduced fertility (as indicated by a decrease in pupal number) (Fig. 2A and fig. S1) in flies compared to normal conditions (25 °C). BG significantly suppressed the influence induced by heat stress on survival in male flies (Fig. 1B) and fertility in female flies (Fig. 2B). A similar trend was observed in the fertility assay using heat-stressed male flies, although the effect was not statistically significant (fig. S1).

**Fig. 1.**
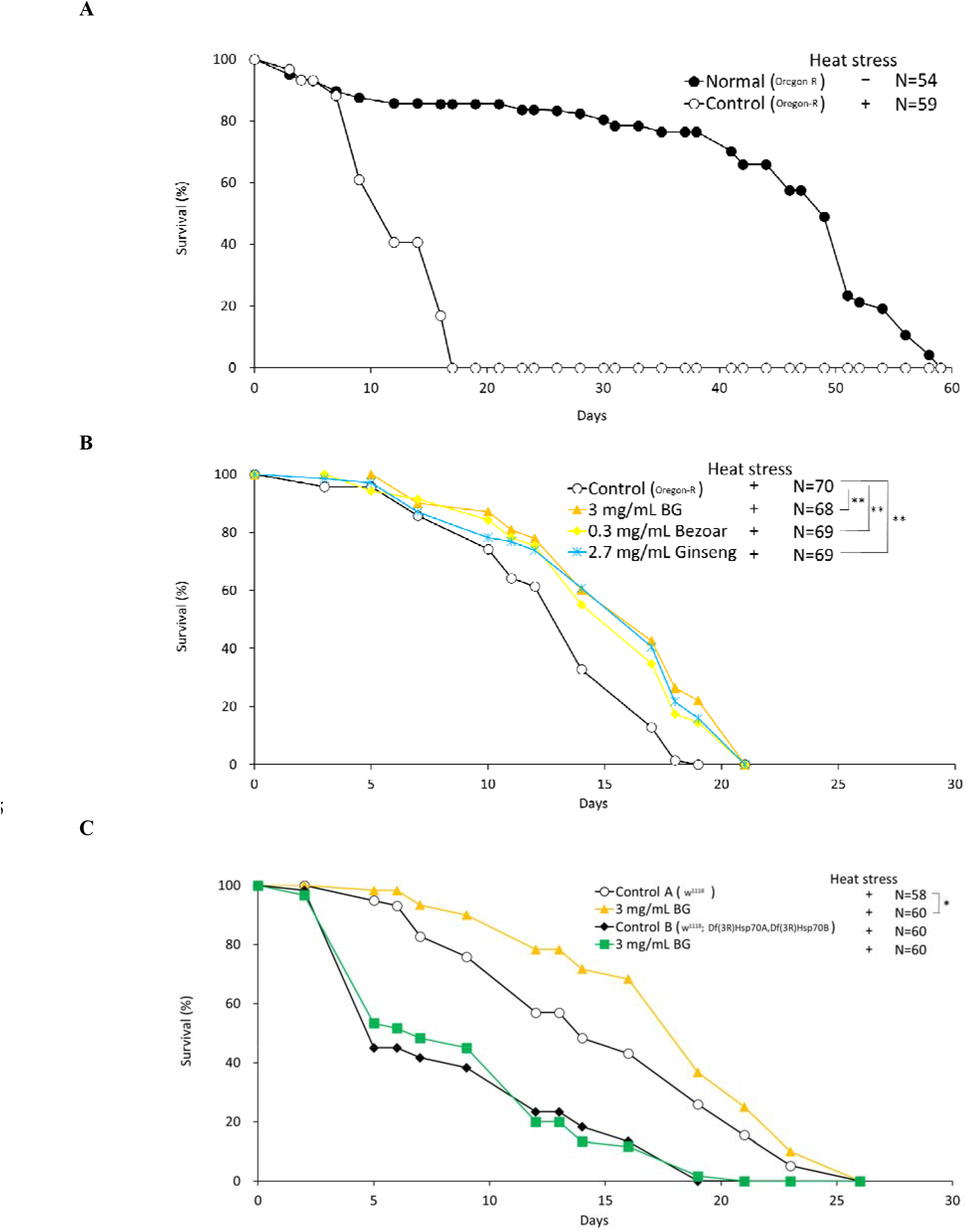
Effects of BG, bezoar and ginseng on the survival in heat-stressed *Drosophila*. (A) Heat stress caused the shortening of survival. (B) BG, bezoar and ginseng suppressed the shortening of the survival. (C) The survival extension induced by BG was not observed in HSP70-null flies. **: P<0.01, *: P<0.05 as compared with the control group using the log-rank test. Bezoar: Oriental bezoar, Ginseng: 2:1 mixture of ginseng and ginseng water extract (dry)

**Fig. 2.**
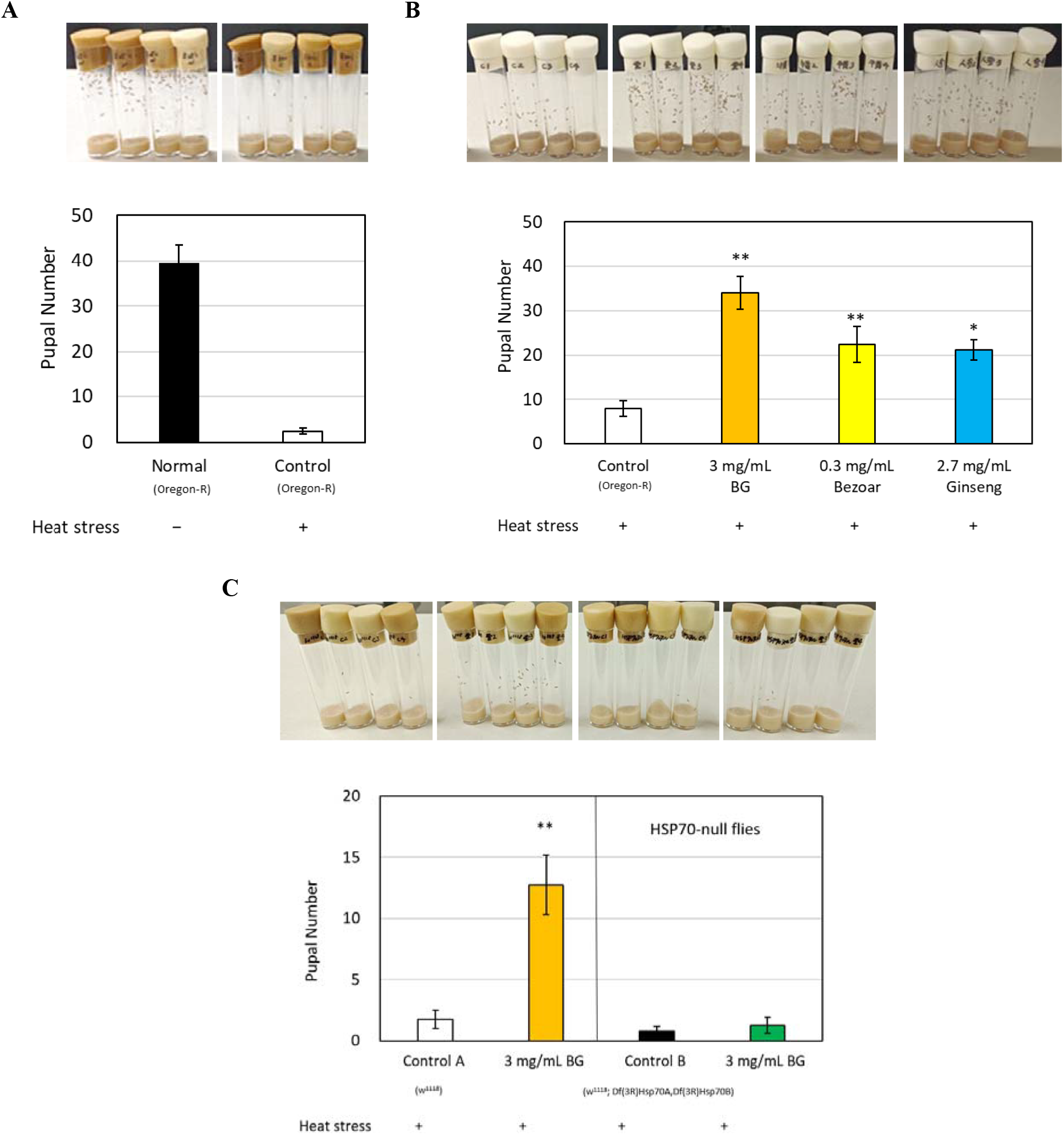
Effects of BG, bezoar and ginseng on the pupal number in heat-stressed female *Drosophila*. (A) Heat stress caused the decrease of the pupal number. (B) BG, bezoar and ginseng suppressed the decrease of the pupal number. (C) The increase of the pupal number induced by BG was not observed in HSP70-null flies. **: P<0.01, *: P<0.05 as compared with the control group using Dunnett’s test or t-test. Data are expressed as the mean ±S.E. (N=7-8). Bezoar: Oriental bezoar, Ginseng: 2:1 mixture of ginseng and ginseng water extract (dry)

### Bezoar and ginseng alone also suppressed shortening of survival and reduction of fertility in heat-stressed Drosophila

BG consists of bezoar and ginseng (2:1 mixture of ginseng and ginseng water extract (dry)). Bezoar and ginseng alone also suppressed shortened survival and reduced fertility in heat-stressed flies; however, their effects were less effective than BG, particularly in the fertility assay (Fig. 1B and 2B).

### HSP70 is essential for BG-mediated protection against heat stress in Drosophila

To investigate the role of HSP70 in the protective effects of BG against heat stress, HSP70-null flies were used. Unlike control flies, HSP70-null flies failed to exhibit BG-mediated protection against heat stress, as BG did not suppress heat stress-induced shortening of survival (Fig. 1C) and reduction of fertility (Fig. 2C). In addition, BG significantly increased HSP70 mRNA expression in heat-stressed flies (fig. S2).

### BG suppressed heat stress-induced cytotoxicity and increased the expression of HSP70 and HSF1 mRNA in HepG2 cells

To evaluate the protective effects of BG against heat stress in humans, the cytotoxicity was examined in heat-stressed HepG2 cells. Heat stress (43 °C) induced cytotoxicity in HepG2 cells compared to normal conditions (37 °C) (Fig. 3A). BG significantly suppressed heat stress-induced cytotoxicity in a dose-dependent manner (Fig. 3B). Next, the effects of BG on the expression of HSP70, HSP90, and their transcription factor HSF1 mRNA were examined. BG significantly increased the expression of HSP70 and HSF1 mRNA, but not that of HSP90 in HepG2 cells (Fig. 4).

**Fig. 3.**
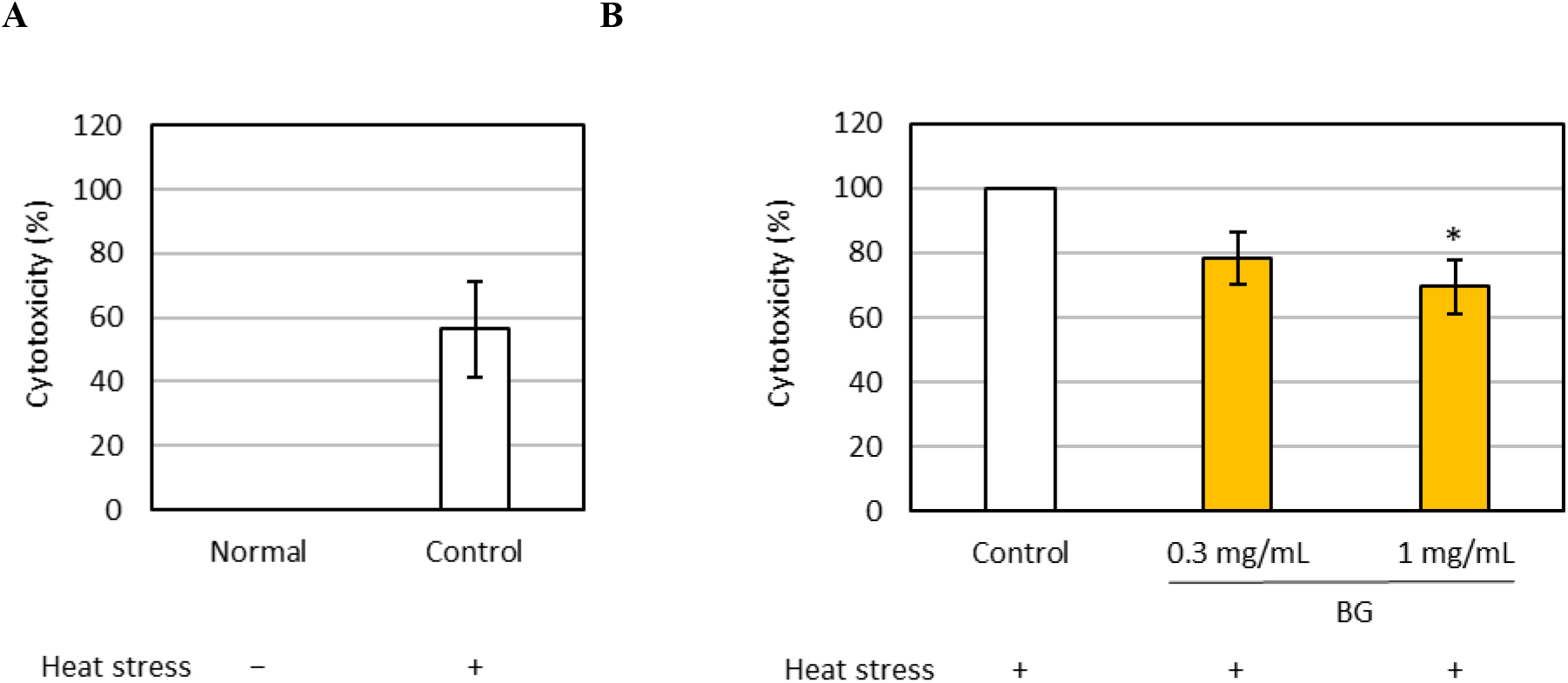
Effect of BG on the cytotoxicity in heat-stressed HepG2 cells. (A) Heat stress at 43 °C for 3.5 h caused severe cytotoxicity. (B) BG suppressed cytotoxicity. *: p<0.05 as compared with the control group using Dunnett’s test. Data are expressed as the mean ± S.E. (N =3).

**Fig. 4.**
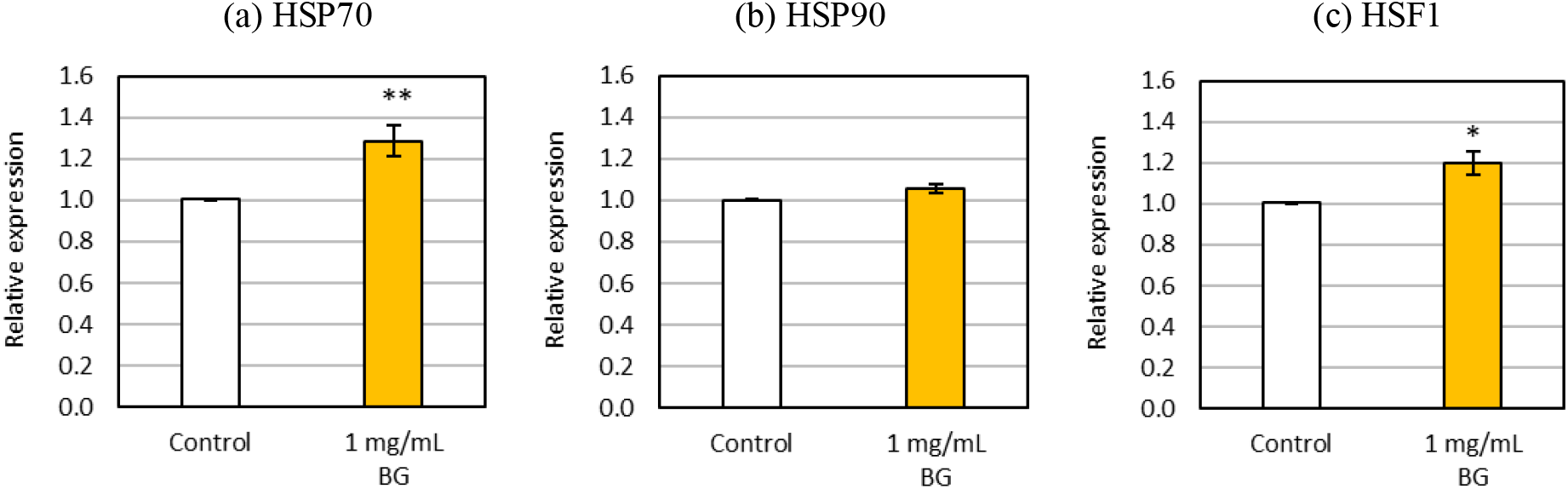
Effect of BG on HSP70, HSP90 and HSF1 mRNA expression in HepG2 cells. BG increased HSP70 and HSF1 mRNA expression. **: P<0.01, *: P<0.05 as compared with the control group using t-test. Data are expressed as the mean ± S.E. (N =4-7).

## Discussion

Heat stress causes a variety of disorders, among which heatstroke is the most severe and life-threatening systemic disease. Recent evidence indicates that heatstroke is not merely a consequence of dehydration, but a complex systemic disorder characterized by endothelial damage, systemic inflammation, dysregulated coagulation/fibrinolysis, and widespread cellular damage (*16*). Although intensive care and cooling therapies are currently used for heatstroke, it remains associated with high morbidity and mortality, and effective pharmacological treatments are still limited (*2, 3*). We found that BG showed protective effects against heat stress in both *Drosophila* and the human hepatic cell line HepG2. BG significantly suppressed heat stress-induced shortening of lifespan, reduction of fertility, and cytotoxicity, suggesting that BG has broad protective effects against heat stress-induced damage. This suggests that BG has potential as a therapeutic agent for heat stress-induced disorders, including heatstroke.

Heat stress induces oxidative stress and protein denaturation, leading to cellular damage, tissue dysfunction, and impaired physiological function (*1*). Among the cellular defense mechanisms against heat stress, HSP70 is one of the most important molecular chaperones that prevents protein aggregation and promotes refolding of damaged proteins (*6, 7*). HSP70 is widely recognized as a promising therapeutic target for heat stress-related disorders (*17*). Geranylgeranylacetone (GGA), an anti-ulcer drug and a well-known HSP70 inducer, improves survival in experimental heatstroke and protects cultured neurons from heat injury by inducing HSP70 (*18, 19*). BG increased HSP70 mRNA expression in both heat-stressed flies and HepG2 cells. Moreover, the protective effects of BG were abolished in HSP70-null flies. These results indicate that HSP70 upregulation is essential for the anti-heat-stress effects of BG. In addition, BG increased HSF1 mRNA expression in HepG2 cells. HSF1 is a master transcription factor regulating HSP70 expression under stress conditions (*11*). Interestingly, GGA induces HSP70 by disrupting the inactive HSP70–HSF1 complex and thereby activating HSF1 (*18*), whereas BG appears to induce HSP70 through transcriptional upregulation of HSF1. These results suggest that BG protects against heat stress-induced damage via the HSF1/HSP70 pathway (Fig. 5). Oriental bezoar and ginseng alone also showed protective effects in heat stress models.

**Fig. 5.**
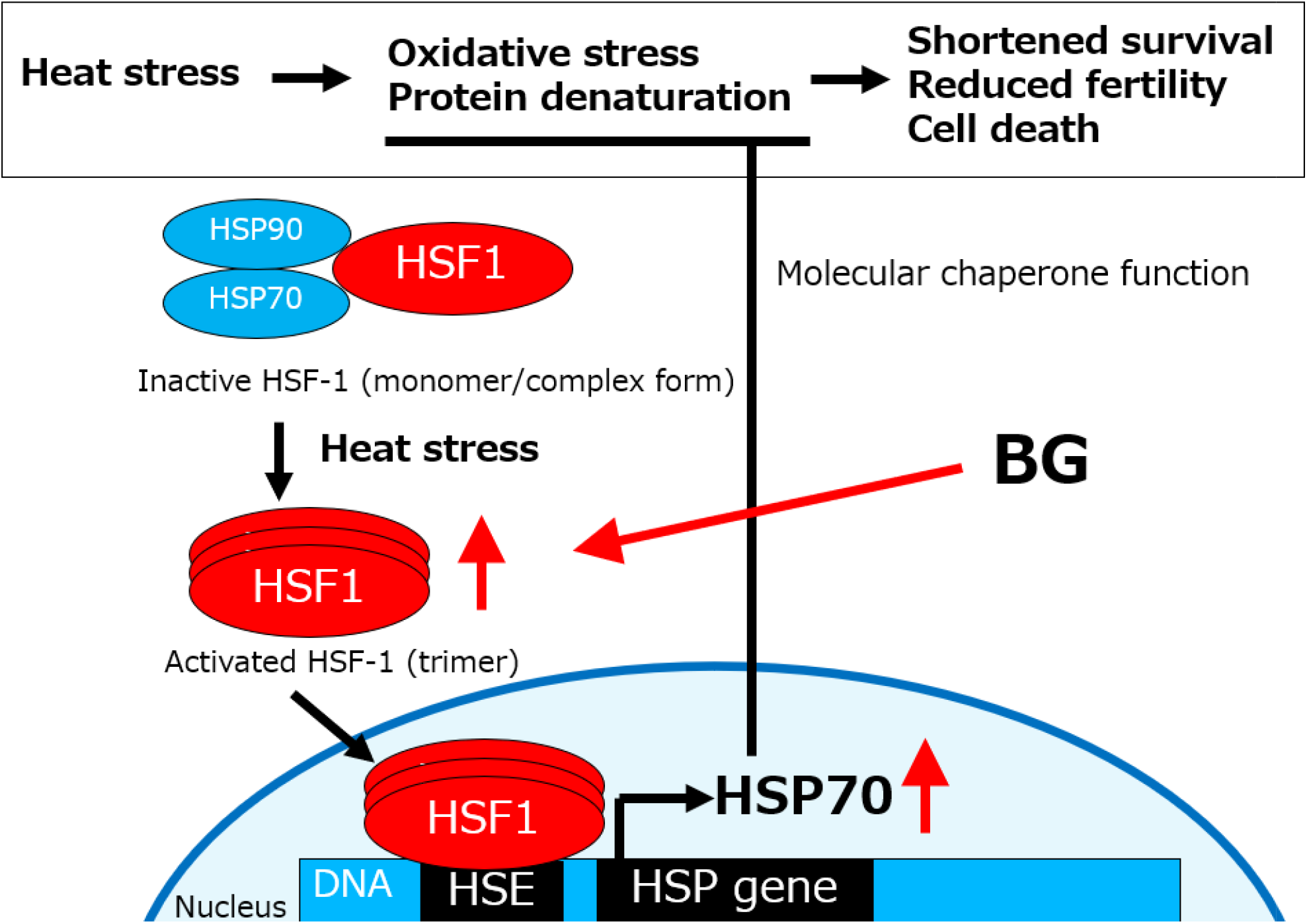
Graphical concepts. Heat stress causes a shortened survival, reduced fertility, and cell death through mechanisms such as oxidative stress and protein denaturation. BG suppresses these adverse effects via HSF1/HSP70 pathway.

Oriental bezoar has traditionally been applied for the treatment of heatstroke (*12*), but its basic pharmacological report has not been found yet. Ginseng has been reported to ameliorate heat stress-induced reproductive dysfunction in Japanese quails (*14*), its molecular evidence remains limited. The present findings show molecular evidence that both oriental bezoar and ginseng possess protective effects against heat stress.

BG exhibited more potent protective effects against the heat-stressed *Drosophila* model than the individual effect of its ingredients, oriental bezoar or ginseng, especially in the fertility assay. These results suggest that the combination of oriental bezoar and ginseng may exert additive or synergistic effects. The Chinese classical medical text Leigong Yaodui describes that “ginseng serves as an assistant to oriental bezoar” (*15*), implying that ginseng and oriental bezoar may act additively or synergistically. The present findings experimentally support this traditional concept and provide molecular evidence for the beneficial combination of oriental bezoar and ginseng.

Several limitations of this study should be acknowledged. Although the involvement of HSP70 in the heat stress protection of BG was supported by the increased HSP70 mRNA expression and the loss of protection in HSP70-null flies, the involvement of the upstream regulator HSF1 was evaluated only at the mRNA level. Further studies examining HSF1 expression and activation are needed to clarify the role of HSF1 in BG-induced HSP70 protein expression. Moreover, the heat stress protection of BG is unlikely to be mediated exclusively through the HSF1/HSP70 pathway. Multiple cellular defense systems against heat stress exist in addition to the HSF1/HSP70 pathway, including the antioxidant Nrf2–ARE, autophagy, and regulation of inflammatory signaling (*20, 21*). BG has shown antioxidant effects *in vitro* and protective effects against ischemia-reperfusion injury *in vivo* (*22, 23*). Further studies are needed to determine whether these pathways contribute to the heat stress protection of BG.

In conclusion, we showed that BG protected against heat stress-induced cellular and organismal damage via the HSF1/HSP70 pathway (Fig. 5). These findings provide molecular evidence supporting the traditional use of oriental bezoar and ginseng for heat stress-induced disorders and suggest that BG has great potential to protect cells and organisms from heat stress-induced disorders, including heatstroke.

Part of this study was presented at the conference (*24–26*).

## Materials and Methods

### Reagents

A Japanese natural drug containing bezoar and ginseng (BG) (Kyushin Pharmaceutical, Tokyo, Japan) contains 100 mg of oriental bezoar, 600 mg of ginseng, and 300 mg of ginseng water extract (dry) (equivalent to 900 mg of ginseng). Oriental bezoar was derived from Brazil, and ginseng was derived from China. For *in vivo* studies, BG was suspended in distilled water and added to the *Drosophila* medium. For *in vitro* studies, BG was suspended in 10% dimethyl sulfoxide and extracted using an ultrasonic generator (Model 2510, Branson Ultrasonics, Branson, CT, USA) for 30 min at 25°C. The extract was sterilized by filtration using a 0.2 µm membrane filter (ADVANTEC, Tokyo, Japan) and added to the culture medium.

Fetal bovine serum, TRIzol^®^, 2.5% trypsin (Thermo Fisher Scientific, Waltham, MA, USA), penicillin-streptomycin (Sigma-Aldrich, St. Louis, MO, USA), isopropyl alcohol (Yoneyama Yakuhin Kogyo, Osaka, Japan), chloroform, dimethyl sulfoxide, ethanol, RPMI 1640 medium (FUJIFILM Wako Pure Chemicals, Osaka, Japan), distilled water (Otsuka Pharmaceuticals, Tokyo, Japan), PrimeScript™ RT Master Mix, TB Green^®^ Premix Ex Taq™ II (Takara Bio, Shiga, Japan) and Cytotoxicity LDH Assay Kit-WST (Dojin, Kumamoto, Japan) were used. PCR primers for HSP70 (human; forward 5′-CCTGGAGTCCTACGCCTTCAAC-3′ and reverse 5′-CTTGACACTTGTCCAGCACCTTC-3′, *Drosophila*; forward 5′-TCGAGATTGACGCACTGTTTG-3′ and reverse 5′-ATCCATCTTGGCATCGTTGAG-3′), HSP90 (human; forward 5′-CAGTACGCTTGGGAGTCCTCA-3′ and reverse 5′-TTTGTTCCACGACCCATAGGTTC-3′), HSF1 (human; forward 5′-TTCGACCAGGGCCAGTTTG-3′ and reverse 5′-CTGCTCGATGTGGACCACTTTC-3′), β-actin (human; forward 5′-TGGCACCCAGCACAATGAA-3′ and reverse 5′-CTAAGTCATAGTCCGCCTAGAAGCA-3′), and RpL32 (*Drosophila*; forward 5′-GCCACCAGTCGGATCGATA-3′ and reverse 5′-GTGCGCTTGTTCGATCCGTA-3′) (Takara Bio) were used.

### Fly stocks

The wild-type strain Oregon R, Hsp70-null strain w^1118^, Df(3R)Hsp70A, Df(3R)Hsp70B, and Hsp70-null genetic background strain w^1118^ were obtained from the Bloomington Drosophila Stock Center (Bloomington, IN, USA). The flies were maintained as previously described (*27*). The flies were reared in vials of a standard medium at 25°C under light-dark (LD) 12:12 conditions. The standard medium consisted of 8% corn meal, 5% glucose, 5% dry yeast extract, 0.64% agar, 0.5% propionic acid and 0.5% butyl p-hydroxybenzoate (*28*). Each drug was mixed into the standard medium.

### Survival assay

Male flies were collected upon eclosion and divided into multiple groups. The flies were housed at a density of 10 flies in a vial containing the medium mixed with the drug and exposed to heat stress at 30°C. The flies were transferred to another vial with fresh medium every three or four days.

The flies were observed during this experiment, and the survival rate of each group was calculated three or four times a week. The flies were considered dead when they did not move despite agitation (*29, 30*).

### Fertility assay

The fertility assay was performed as previously described (*31*), with minor modifications. Female or male flies (1–3 days old) were placed in vials containing the medium mixed with the drug and exposed to heat stress at 30°C for 4 days. These flies were then transferred to vials containing 4-10 days old flies of the opposite sex (3 males and 3 females/vial) and allowed to mate at 25°C for 24 h. Seven days after fly removal, the pupal number formed in the vial was counted. Additionally, HSP70 mRNA expression was measured by RT-PCR from the flies following heat stress at 30°C for 4 days.

### Cell culture

HepG2 cells were purchased from RIKEN BioResource Research Center (Tsukuba, Japan) and maintained in RPMI 1640 with 10% FBS, 100 U/mL penicillin, and 100 μg/mL streptomycin at 37°C in 5% CO_2_.

### Cytotoxicity assay

HepG2 cells were cultured in 96-well microplates (Corning, Corning, NY, USA) at a density of 1 × 10^4^ cells/well/200 µL for 48 h, then the medium was replaced with a new medium containing each drug, and the cells were pre-incubated for 2 h. The cells were exposed to heat stress at 43 °C for 3.5 h, then the drug was washed out, and the cells were incubated at 37 °C for 48 h. Cytotoxicity was calculated by LDH assay using the Cytotoxicity LDH Assay Kit-WST. The absorbance of the formazan dye in the well at 490 nm was measured using a microplate reader (Synergy H1, BioTec, Winooski, VT, USA).

### mRNA expression analysis

HepG2 cells were cultured in 48-well microplates (Corning) at a density of 2 × 10^4^ cells/well/200 µL for 48 h, then the medium was replaced with a new medium containing each drug and the cells were incubated for 2 h. After removing the medium, 200 µL of TRIzol^®^ was added to the cells. The cells were scraped to disrupt cell membranes, left for 5 min at room temperature, and collected in a microtube. Then, 40 µL of chloroform was added, and the mixture was shaken vigorously and stirred. After 5 min, each sample was centrifuged at 12,000 × g and 4°C for 15 min. The supernatant (50 µL) was transferred to a new microtube. Then, 70 µL of isopropyl alcohol was added, and the mixture was stirred. After 10 min, each sample was centrifuged at 12,000 × g and 4°C for 10 min. After removing the supernatant, 200 µL of 75% ethanol was added, and the mixture was stirred. Each sample was centrifuged at 7,500 × g and 4°C for 5 min, and the supernatant was removed. The precipitates were dissolved in 40 µL of distilled water. A 20 µL reverse transcription reaction mixture consisting of 2 µL of 5 × PrimeScript™ RT Master Mix, 0.5 µg of total RNA, and distilled water was applied to the reverse transcription reaction at 37°C for 15 min and 85°C for 5 s. A 25 µL real-time PCR mixture consisting of 12.5 µL of TB Green^®^ Premix Ex Taq™ II, 0.2 µL of 50 µM forward primer, 0.2 µL of 50 µM reverse primer, less than 100 ng of 2.0 µL template DNA, and 10.1 µL of distilled water was applied to the PCR (Thermal Cycler Dice Real Time System II, Takara Bio) at 95°C for 30 s, and 35 cycles of two-step PCR were performed at 95°C for 5 s and 60°C for 30 s. The mRNA levels were quantified by calculating ΔCt and ΔΔCt values.

### Statistical analysis

The results were expressed as means ± standard error. Significant differences in data were estimated using the log-rank test, Student’s t-test or one-way analysis of variance followed by Dunnett’s multiple-comparison test. The differences with P < 0.05 were considered statistically significant.

## Author contributions

Conceptualization: EI, TT, YS, HK, NI

Methodology: EI, TT, YS, HK, NI

Investigation:EI, TT

Visualization: EI, TT

Funding acquisition: NI

Project administration: NI

Supervision: HK, YS, NI

Writing – original draft: EI, TT

Writing – review & editing: HK, YS, NI

## Data, code, and materials availability

All data are available in the main text.

---

**Fig. S1.**
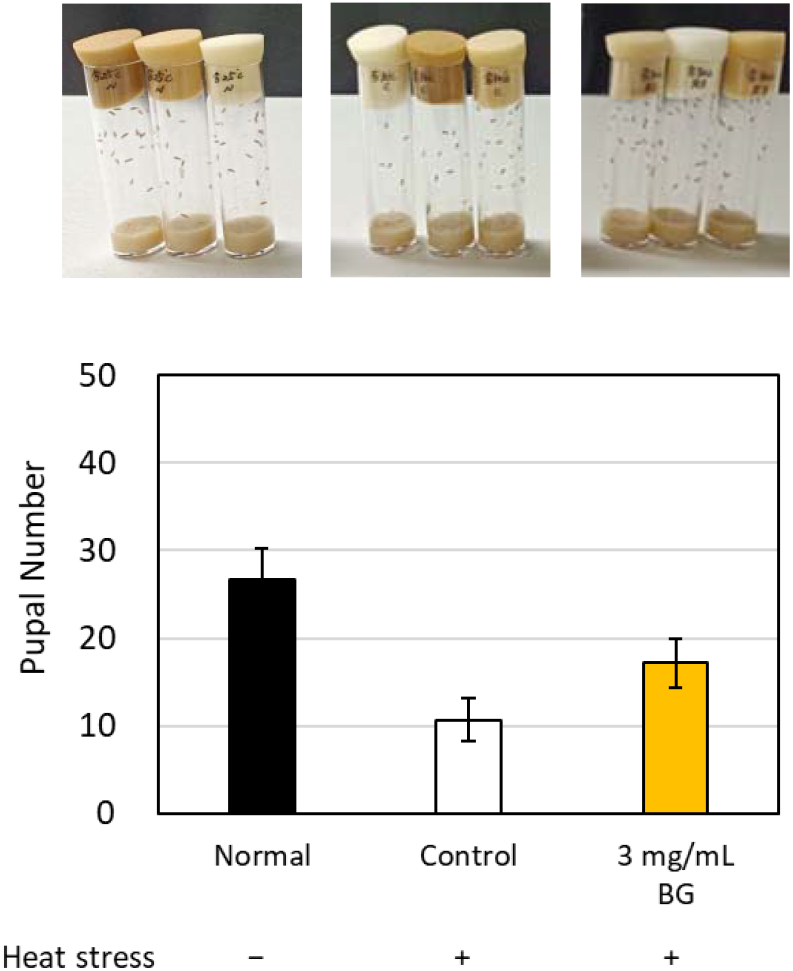
Effect of BG on the pupal number in the fertility assay using heat-stressed male *Drosophila*. BG did not significantly suppress the decrease of pupal number in the fertility assay using heat-stressed male fly. Data are expressed as the mean ±S.E. (N=7-8).

**Fig. S2.**
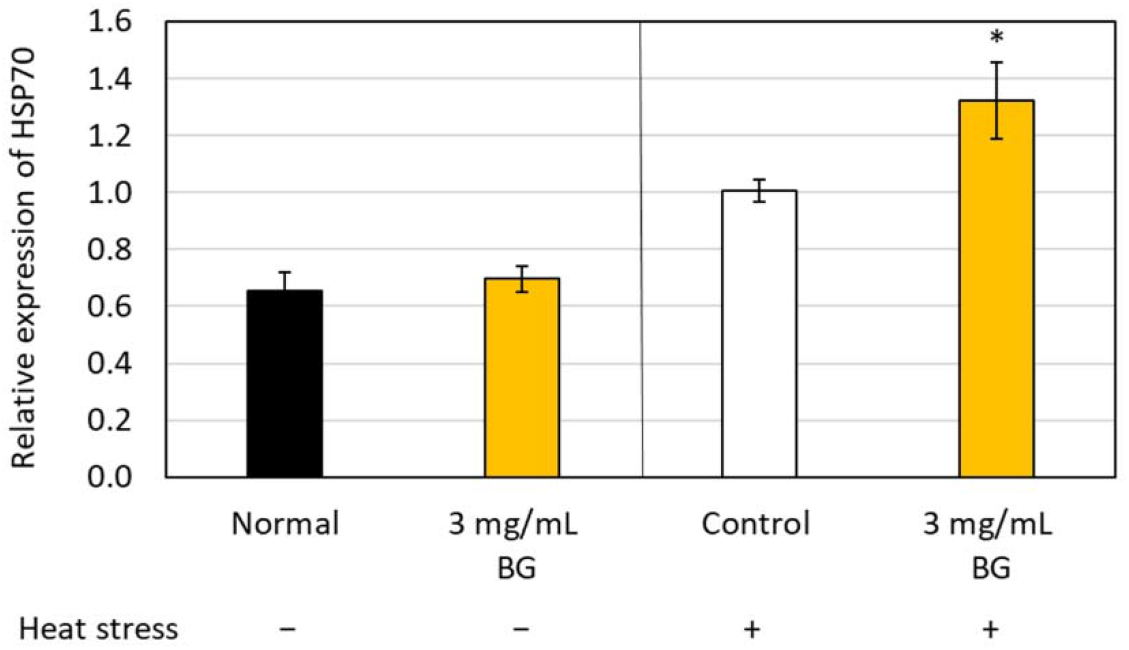
Effect of BG on HSP70 mRNA expression in heat-stressed female *Drosophila*. BG increased HSP70 mRNA expression in heat-stressed female flies. *: p<0.05 as compared with the control group using t-test. Data are expressed as the mean ±S.E. (n=7-8).

